# Development of Human Neuroblastomas in Mouse-Human Neural Crest Chimeras

**DOI:** 10.1101/523795

**Authors:** Malkiel A. Cohen, Shupei Zhang, Satyaki Sengupta, Haiting Ma, Brendan Horton, George W. Bell, Rani E. George, Stefani Spranger, Rudolf Jaenisch

## Abstract

Neuroblastoma (NB), derived from the neural crest (NC), is the most common pediatric extracranial solid tumor. Here we establish a platform that allows studying human NBs in mouse-human NC chimeras. Chimeric mice were produced by injecting human NC cells carrying NB relevant oncogenes *in-utero* into gastrulating mouse embryos. The mice developed tumors composed of a heterogenous cell population that closely resembled that seen in primary NBs of patients but were significantly different from homogenous tumors formed in xenotransplantation models. The human tumors emerged in immunocompetent hosts and were extensively infiltrated by mouse cytotoxic T cells reflecting a vigorous host anti-tumor immune response. However, the tumors blunted the immune response by inducing infiltration of regulatory T cells and expression of immune checkpoints similar to escape mechanisms seen in human cancer patients. Thus, this experimental platform allows studying human tumor initiation, progression, manifestation and tumor – immune-system interactions in an animal model system.

## Introduction

Based on the discovery of cancer immune checkpoints and the success of checkpoint inhibitors, the new generation of cancer immunotherapies have resulted in remarkable advances in cancer treatment. However, the fraction of patients who respond to immune therapy is generally around 20% for the most common solid tumors (Ribas and Wolchock, 2018) calling for model systems that would facilitate studying the parameters that allow tumors to escape immune inhibition. Human cancers are commonly studied in xenotransplantation models involving the injection of human cancer cells or primary tumors into immunocompromised mice. While these models have yielded a wealth of information on the biology of human cancer as well as on therapeutic strategies, they pose several limitations. Because only end stage tumor cells, often adapted to cell growth in culture, are transplanted into host animals that are immune deficient, these xenotransplantation models do not allow investigating the initiation of the tumor and the anti-tumor immune reaction or tumor immune evasion. An alternative experimental approach for studying the anti-cancer immune response *in vivo* is tumor induction in transgenic mice. However, species specific differences between human and mouse may make it problematic to relate results obtained in transgenic mouse models to human cancer.

Interspecies chimeras represent a promising experimental system for studying human development and disease and may provide the most physiologically relevant environment to study human disease in an *in vivo* context by overcoming some of the limitation of conventional xenotransplantation animal models (Wu et al., 2016; Suchy and Nakauchi 2017, Soldner and Jaenisch, 2018). Both, pluripotent and committed stem cells have been used as donor cells to generate interspecies chimeras. Injection of pluripotent rat stem cells (PSCs) into mouse blastocysts resulted in chimeric mice with rat donor cell contributing to all tissues (Kobayashi et al., 2010). In contrast, human PSCs introduced into mouse blastocysts resulted in very low if any functional incorporation of the human donor cells into the host embryo (Gafni et al., 2013, Theunissen et al., 2016, Wu et al., 2017, Yang et al., 2017) with no postnatal chimeras having been generated.

Interspecies postnatal chimeras have been produced by introducing multipotent, lineage-restricted stem cells into post-implantation mouse embryos or neonates. For example, human glial progenitors injected into the neonatal mouse brain integrated into the host brain were shown to enhance synaptic plasticity and learning in the chimeric mice (Han et al., 2013; Windrem et al., 2017). Similarly, hPSCs derived neurons transplanted into Alzheimer’s disease (AD) mouse model display signs of neurodegeneration presented by cell death and classical pathological features typically observed in AD patients (Espuny-Camacho et al., 2017). We have, based on previous *in-utero* manipulations of mouse embryos (Jaenisch, 1985, Huszar et al., 1991), generated mouse-human neural crest chimeras (Cohen et al., 2016). Neural crest cells (NCCs) are multipotential, emerge from the neural tube at gastrulation and generate a wide variety of lineages including peripheral neurons, enteric neurons, Schwann cells, melanocytes and cells of adrenal medulla (Bronner and LeDouarin, 2012). Neural crest (NC) related deficiencies are the cause of multiple human diseases and the term “neurocristopathies” has been proposed to denote syndromes or tumors involving NCCs. Neural crest derived malignancies include cancers such as melanoma, neuroblastoma and neurofibromatosis (Vega-Lopez et al., 2018) and the use of hPSCs-derived NCCs for modeling human NC diseases is an attractive *in vitro* experimental approach (Fattahi et al., 2016, Huang et al., 2016). We showed that hPSCs-derived NCC, when injected into the gastrulating mouse embryo, migrated along the dorso-lateral migration route contributing to the pigment system of the mouse, suggesting that this platform may be used to model neurocristopathies *in vivo (*Cohen et al., 2016).

The goal of this work was to model human neuroblastoma (NB), one of the most common extracranial childhood tumors. NB is an embryonal cancer derived from the developing neural crest, with tumors arising in sympathetic ganglia and adrenal medulla (Cheung and Dyer, 2013). We have generated hPSCs and hNCCs carrying the Doxycycline (Dox) inducible NB-relevant oncogenes *MYCN* and *ALK*^*F1174*^ (Berry et al., 2012, Schulte et al., 2013). When the hNCCs were injected into mouse embryos, tumors resembling primary human NBs were formed. Most significantly, the tumors developed in immunocompetent host mice allowing the analysis of human tumor-immune system interaction and immune escape mechanisms in an experimental animal system. We found that the host mounts an anti-tumor immune response that is blunted by tumor induced immune inhibitory mechanisms. This chimeric system facilitates the study of critical interactions between NBs and the immune system, in contrast to conventional xenotransplantation assays that rely on immunocompromised host animals and thus preclude meaningful study of the immune response to cancer development.

## Results

### Generation of human pluripotent cells carrying conditional oncogenes

Previous data have shown that human NC cells injected into E 8.5 embryos migrate along the dorso-lateral migration route and can contribute to the host pigment system in postnatal mice (Cohen et al., 2016). In these studies, we did not investigate whether the human donor cells can also migrate along the medial NC migration route and contribute to internal structures (Trainor, 2005). To this end, hNCCs were microinjected into E8.5 mouse embryos and the embryos were analyzed 5 to 6 days post-injection (**Figure S1 A**). eGFP positive cells were found in dorsal root ganglia (DRG) and in the Trigeminal ganglion (**Figures S1 B-C**), and immunostaining for Peripherin, a PNS marker, confirmed proper integration of human cells within the host DRGs (**Figure S1 B,** right). To assess hNCC contribution into postnatal mice, we individually tested DRGs of NC chimeric mice (confirmed by coat-color contribution) for human contribution by qPCR (Cohen at al., 2016). Human contribution was found in about 10% of tested DRGs of chimeric mice (**Figure S1 D**), indicating that the hNCCs can contribute, in addition to the pigment lineage, also to the peripheral nervous system. However, we failed to detect human NC donor cells in structures of the autonomic nervous system such as in the intestine or the adrenal gland. Because colonization of these structures would require a long migration of the donor cells through the developing embryo, our failure to detect the presence of donor cells at the autonomic nervous system may have been due to low contribution. As shown below, providing the donor NC cells with a proliferative advantage such as expression of NC-tumor relevant oncogenes enhanced detection of rare donor cells contributing to the autonomic nervous system.

We generated hPSCs, which conditionally overexpress MYCN and *ALK*^*F1174L*^ to model human neuroblastoma. MYCN is frequently amplified in high-risk NB and correlates with an aggressive phenotype and poor prognosis (Brodeur et al., 1984, Seeger et al., 1985). Several mutations in the anaplastic lymphoma kinase (*ALK*) gene are involved in the development of sporadic and familial neuroblastoma (Chen et al., 2008, George et al., 2008, Janoueix-Lerosey et al., 2008, Mossé et al., 2008). One of the common mutations in the *ALK* gene results in a cytosine-to-adenine change in exon 23 and a phenylalanine-to-leucine substitution (F1174L) within the kinase domain, causing abnormal proliferation of neuroblasts. In mouse model systems overexpression of both *Mycn* and the *Alk*^*F1174L*^ mutation was found to be sufficient to transform mouse NCCs *in vitro* and to induce NBs *in vivo (*Berry et al., 2012, Schulte et al., 2013). To conditionally overexpress MYCN and ALK^F1174L^ in human NCCs, we transduced the Dox inducible lentiviral constructs FUW-TetO-*ALK*^*F1174L*^-t2A-*tdTomato* and FUW-TetO-*MYCN*-t2A-*NeoR (***Figure S2 E)** into two hPSC lines (WIBER#3 hESCs and AA#1 hiPSCs; Cohen et al., 2016). To facilitate tracing of the cells after injection, we used hiPSCs which harbored eGFP reporter into the *AAVS1* locus, and inserted Luciferase and Lacz reporter constructs into the *AAVS1* locus of hESCs (**Figures S2 A-D;** Hockemeyer et al., 2009). Using established protocols, the hPSCs were differentiated into hNCCs (Cohen at al., 2016) and tested for Dox dependent activation of the oncogenes and the effect on hNCCs growth. Addition of Dox to the hNCCs culture medium activated the oncogenes, as indicated by the expression of the tdTomato fluorescence reporter, by RNA and protein expression analyses (**Figures 1A-D**). **Figures 1D-F** show that Dox treatment induced the ALK phosphorylation and the down-stream MAPK/ERK signaling pathway, increased cell proliferation and colony formation in soft agar. To test whether oncogene activation would promote tumor formation, the hNCCs were injected into immunocompromised mice. **Figure 1G** shows that Dox dependent expression of the two oncogenes was sufficient to drive tumor formation in immune compromised mice.

**Figure 1.**
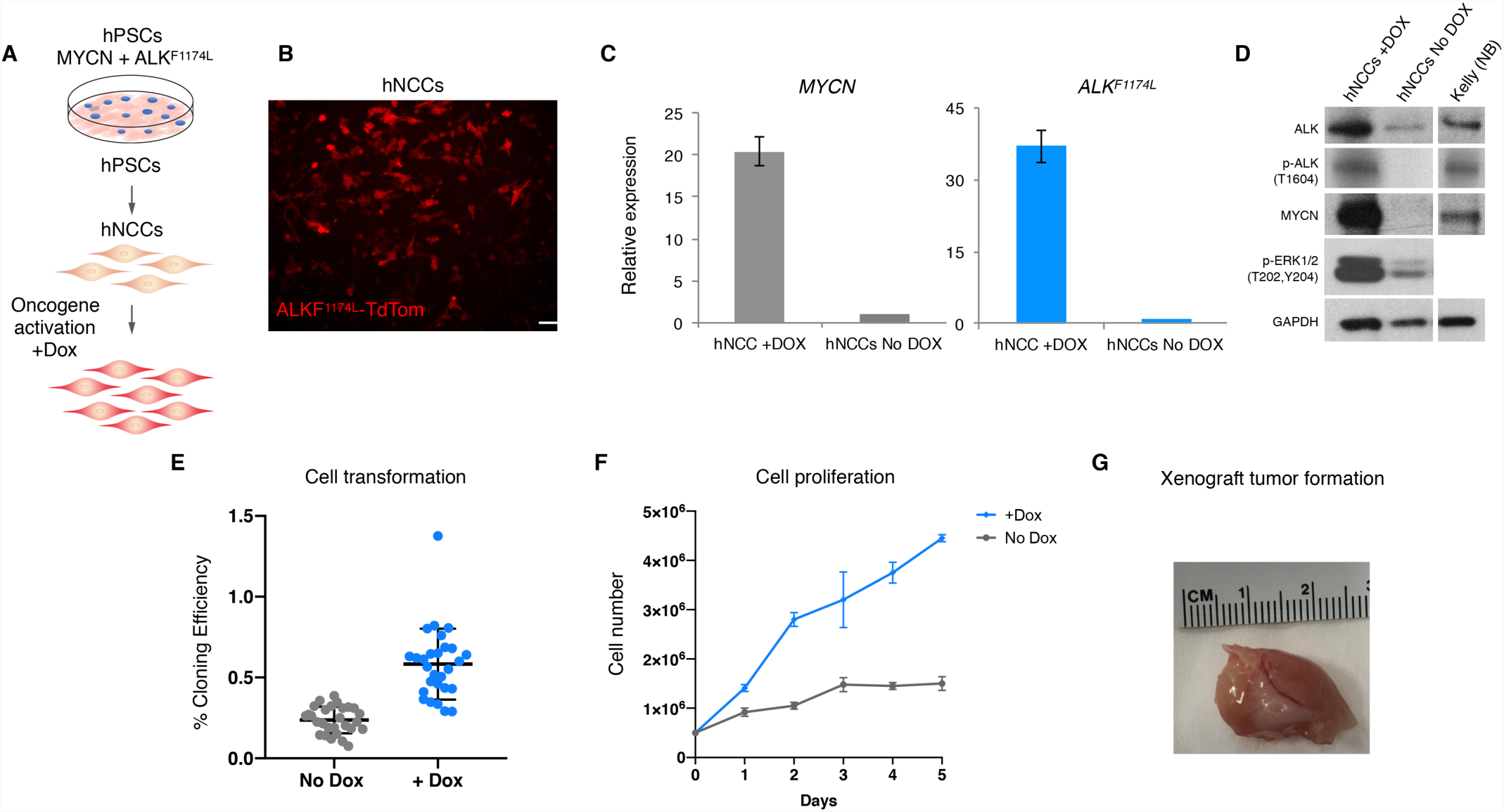
Generating hPSCs Derived Human Neural Crest Prone To Generate NB. (**A**) A schematic representation of the experiment. Viral transgenes of *FUW-TetO-ALK*^*F1174L*^*-t2A-tdTomato* and *FUW-TetO-MYCN-t2A-NeoR* were introduced to hPSCs for the controlled overexpression of ALK^F1174L^ and MYCN. hPSCs were differentiated into hNCCs and oncogenes were activated by Dox. (**B)** By Dox administration, hNCCs expresses ALK^F1174L^ along with tdTomato fluorescence protein (Scale bar =50µm). (**C**, **D**) Oncogenes activation was monitored by gene expression using qRT-PCR and at the protein level by Western blot. Dox treatment induced the ALK phosphorylation and the down-stream MAPK/ERK signaling pathway (**D**, Kelly NB cell line served as positive control). (**E**-**G**) hNCCs with activated oncogenes show characteristics of oncogenic traits as represented by colony forming assays on soft agar (**E**, n=28), measuring the hNCCs proliferation (**F**), and by xenograft formation following injection into immune-compromised NSG mice (**G**).

### Tumor formation in mouse-human neural crest chimeras

To generate mouse-human NC chimeras, the hNCCs were microinjected *in-utero* into developing mouse embryos at E8.5 as previously described (Jaenisch 1985, Cohen at al., 2016). To activate the oncogenes, Dox was added to the drinking water of the pregnant females at day E9.5-10. (**Figure 2A**). To trace cell migration, we inspected embryos at 6 days after injection and found eGFP^+^ donor cell clumps in E-14.5-15.5 injected embryos suggesting cell proliferation of donor cells induced by oncogene activation (**Figure S3**). We first detected Luciferase activity in injected mice by using non-invasive bioluminescent imaging in 3 to 6 months old mice, consistent with proliferation of the injected oncogenic cells (**Figures 2B-C**). About 20% of the injected mice developed an abdominal mass, mostly from the retroperitoneal space, a typical location seen in NB patients (The total number of mice injected as embryos = 144; See **Figure 2C** and **Figure S4**) with no macroscopic tumors detected in other organs. Tumors designated as CHNB (Chimeric derived Neuroblastoma) were collected between 3 and 15 months of age (Mean = 10.4 ± 3.9 month) and found to express eGFP, confirming human donor cell origin. Moreover, the tumors immunostained for MYCN, and ALK and expressed tdTomato, verifying oncogenes expression in the human tumors (**Figures 2 D-G**). Also, the human tumors were positive for phosphorylated H2AX (γH2AX), a DNA double-strand break (DSB) marker, indicating genome instability, which is typically associated with malignancy (**Figures 2H**). Similar to high-grade NB in patients, these CHNBs were highly proliferating as indicated by the expression of the cell cycle marker Ki67 (**Figures 2I**).

**Figure 2.**
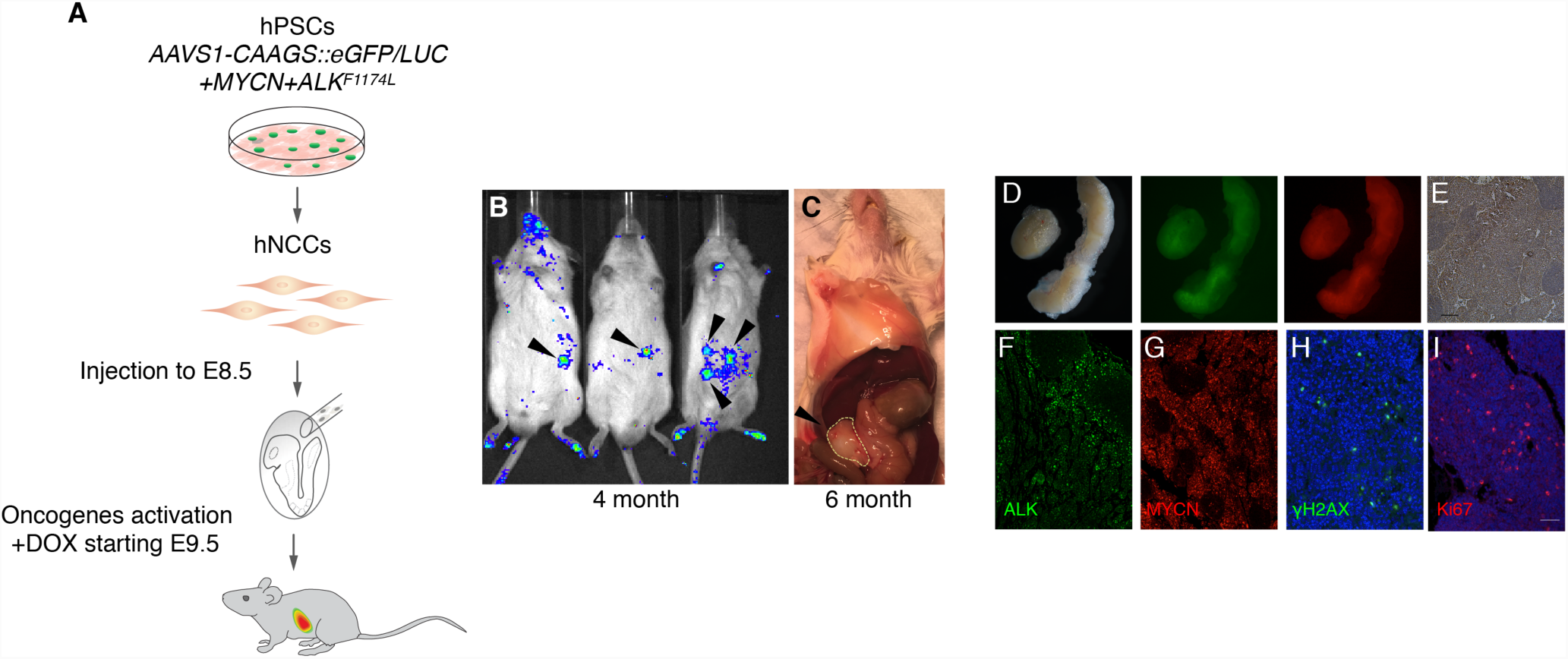
hNCCs Develop Human Tumors in Mouse-Human Neural Crest Chimeras. (**A**) A schematic representation of the experiment. E8.5 mouse embryos injected with hNCCs were found to form human tumors originating from donor hNCCs. (**B**) Non-invasive bioluminescent imaging shows the first indication of human tumor formation at the age of 3 month, and (**C**) macroscopic growth were first detected at the age of 6 month. (**D**) Tumors of chimeric mice dissected at P120 exhibit eGFP and tdTomato fluorescence, indicating their origin from donor hNCCs. (**E**) eGFP expression within tumors was confirmed by IHC (scale bar =100µm). (**F**-**I**) The human tumors were found to express MYCN and ALK expression (**F**-**G**), and the typical cancer hallmarks γH2AX and Ki67 (**H**-**I**), by IF (scale bar =20µm).

Histological examination of haematoxylin-eosin (H&E) stained CHNB tumors revealed typical densely packed primitive neuroblasts with lobular patterns, which is similar to primary tumors from of patients. IHC showed that CHNB human tumors consisted of different cell types expressing a mixture of markers such as Tyrosine Hydroxylase (TH; 46.71 ± 18.29%), Synaptophysin (SYP; 44.45 ± 10.95%), Chromogranin A (CgA; 58.05 ± 16.20%), and Nestin (NES; 37.38 ± 22.69%) in heterogeneous patterns, all of which are typical markers for primary human NBs (**Figure 3A,** middle and right panels and **Figure S5**). In contrast, xenograft-tumors derived from hNCCs overexpressing *ALK*^*F1174L*^ and *MYCN* and injected into immunocompromised mice were homogenous and lacked cells expressing the typical NB markers (**Figure 1G, Figure 3A** left panels, and **Figure S5)**. Gene expression profiling by RNA-Seq revealed a significant correlation in the expression levels of 20 frequently activated NB-associated genes between tumors growing in chimeric mice and their expression seen in human NBs (**Figure 3B**; *p*-value=0.011; Harenza et al., 2017). These results demonstrate that hNCCs expressing *MYCN* and *ALK*^*F1174L*^, when integrated into tissues of the developing mouse embryo, induce the growth of tumors that resemble primary tumors isolated from NB patients but are very different from tumors produced in conventional xenograft models.

**Figure 3.**
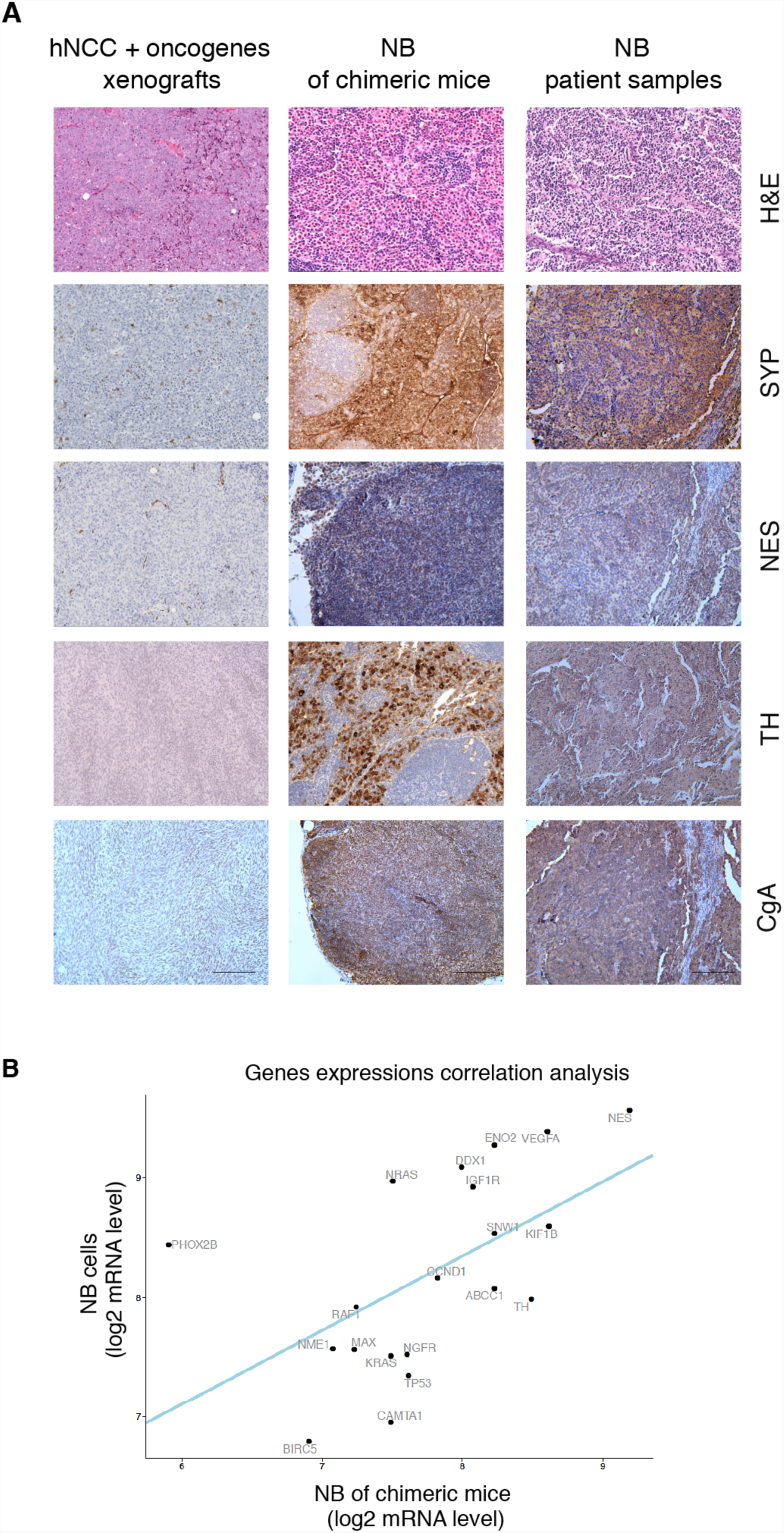
Tumor of Mouse-Human Neural Crest Chimeras Present Phenotypes of Human NB *in Vivo.* (**A**) H&E and IHC comparing assay show that human NB tumors of chimeric mice express the typical NB markers Synaptophysin (SYP), Nestin (NES), Tyrosine hydroxylase (TH) and Chromogranin A (CgA), similarly to their expression found in NB samples of patient. hNCCS, which expressed oncogenes and were subcutaneously injected into immunocompromised mice to form xenograft tumors (left column) did not express these NB markers. (See IHC quantifications in **Supplementary Figure 4**; scale bars =100µm). (**B**) RNA-Seq of CHNB tumor samples (n=4) were separated *in-silico* into human and mouse reads to separate the tumor and hosts’-environment compartments (See material and methods). The analysis of RNA-Seq of the human-gene expression profile revealed that the human tumors in chimeric mice expressed a set of key genes normally associated with NB tumors (*ABCC1, BIRC5, CAMTA1, CCND1, DDX1, ENOS, IGF1R, KIF1B, KRAS, MAX, NES, NGFR, NME1, NRAS, PHOX2B, RAF1, SNW1, TH, TP53* and *VEGFA*) with a significant correlation to expression in NB cell lines (Kelly and SHSY-5Y). linear regression *p*-value =0.011.

### Tumors form in immunocompetent recipient mice

In contrast to conventional xenograft models, the human cells were found to form NBs in immunocompetent hosts with the fraction of NB formation (number of tumors in total number of mice injected as embryos) being similar to the fraction in immunocompromised hosts (**Figure S4**). Growth of human tumors in immunocompetent mice suggests that the hosts were tolerized to the human cells (Xing and Hogquist, 2012). To test this, we investigated the immune-microenvironment of the human NBs that had developed in the immunocompetent mice. Immuno-histological analysis of NBs growing in adult chimeric mice showed strong infiltration by host (murine) CD3^+^ and cytotoxic CD8a^+^ T cells and macrophages consistent with a significant host immune response, and high immune score, similar to patient derived inflamed tumors (Galon et al., 2006; **Figure 4A** and **Figures S6A-C**). Because CD8^+^ cytotoxic T cells are the main effectors in various cancer types triggering tumor cell destruction, we investigated potential mechanisms of immune escape of CHNB tumors. **Figure 4C** shows that the human NBs recruited regulatory T cells (Tregs) stained for FoxP3 and Il2ra consistent with induction of an anti-tumor immune response (**Figure S6D)**. In addition, the tumors activated immune checkpoint signals as indicated by staining for murine specific Tim-3 in host T cells and human specific PD-L1 (a ligand of PD-1) and CD47 expressed in tumor cells (**Figure 4D** and **Figures S6D-E**) suggesting a mechanism for the human tumors to escape immune destruction (Woo et al., 2001, Dong et al., 2002, Jaiswal et al., 2009, Fourcade et al., 2010). Finally, gene expression profiling by RNA-Seq of the host’s murine micro-environment confirmed a gene signature typical for activated T cells and tumor inflammation (**Figures 4E-F**). In contrast, T cell activation within the tumor microenvironment was not seen in a transgenic mouse model of NB (Berry et al., 2012), which lack the infiltration by immune cells (**Figure 4B** and **Figures S6A-C**). Our results suggest that human tumors growing in chimeric mice evade a strong anti-tumor immune response by similar escape mechanisms as seen in primary human tumors.

**Figure 4.**
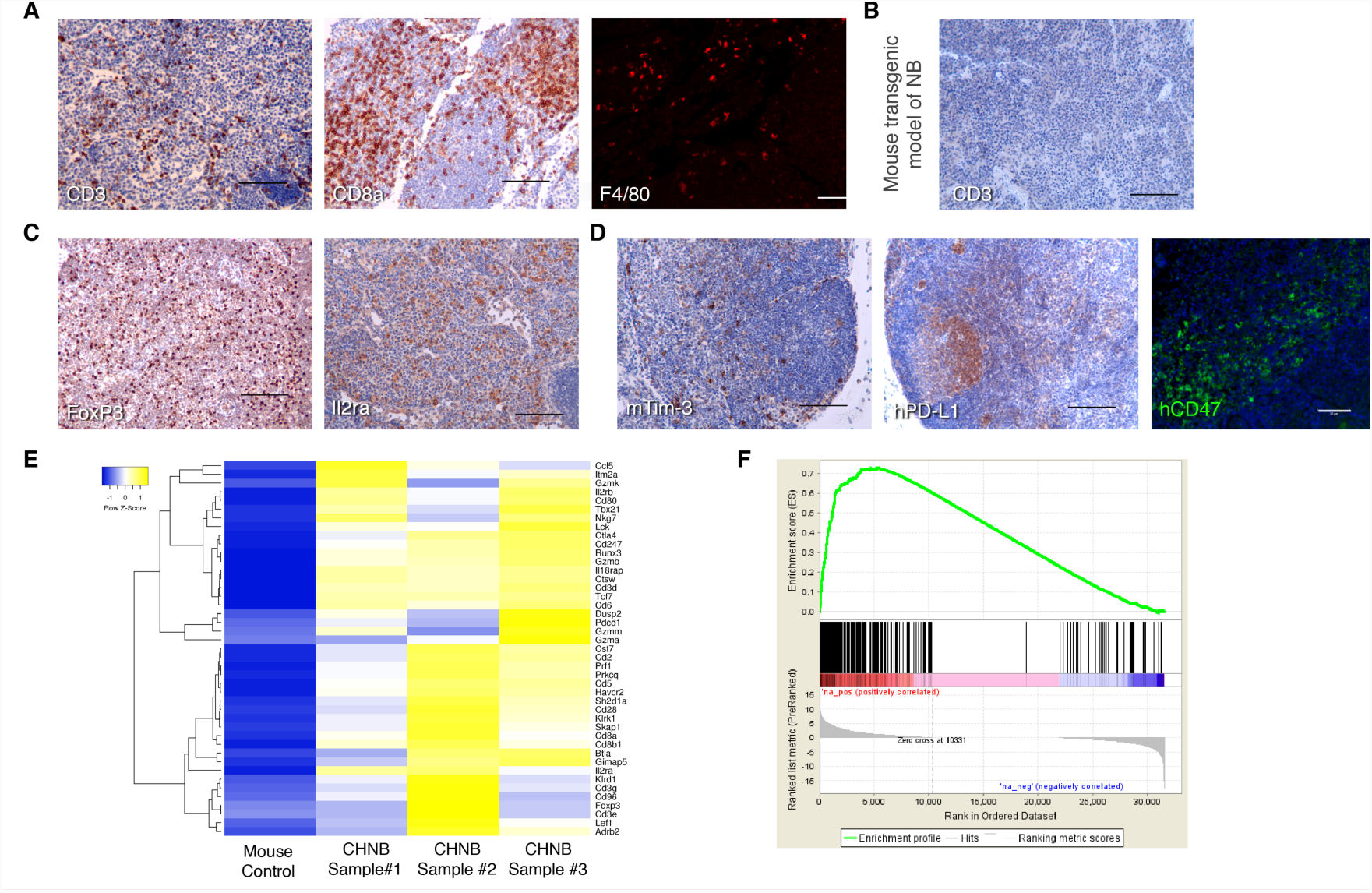
Characterization of the Immune-Microenvoirment of the NB Chimeric Model. **(A)** Host T cell, CTL and macrophage infiltration marked by CD3, CD8a and F4/80 in NB tumors of chimeric mice. (**B**) No CD3^+^ T cells infiltrate was observed in a NB mouse model of transgenic mice (Berry et al., 2012). (**C**) Human NB tumor of chimeric mice shows immune tolerance marked by infiltration of Tregs (staining for FoxP3) activated cells (Il2ra). (**D**) T cell exhaustion was observed in NB chimeric mice, indicated by the expression of immune checkpoint signals (mouse specific Tim-3, human specific PD-L1 and human specific CD47; IHC scale bars =100µm, IF scale bars =50µm). (**E**, **F**) RNA-Seq of CHNB tumor samples were sorted *in-silico* into human and mouse reads (See material and methods). The RNA-Seq of the murine-genes, representing hosts’ tumor microenvironment, show a gene expression signature typical of immune infiltrates and cancer related inflammation response.

### Cytotoxic T cells recognize oncogene-related antigens on human NBs

To investigate the cross talk between immune and tumor cells we isolated T cells from the chimeric mice to test any direct interactions between the host’s immune system and the human tumor cells. Splenocytes of chimeric mice that developed macroscopic CHNB tumors were isolated and co-cultured with the donor hNCCs to test T cell activation as a measure for specific recognitions of the TCR and an inducing epitope/s (**Figure 5**). Co-cultures with hNCCs that did not express the oncogene did not stimulate T cell proliferation, similar to other human cell controls like human dermal fibroblast (expressing the human leukocyte antigen class I; HLA-I), and MYCN-amplified Kelly NB cells (negative for HLA-I, see **Figure S7**). In contrast, when the T cells were co-cultured with hNCCs overexpressing *MYCN* and *ALK*^*F1174L*^ T cell proliferation was highly induced (**Figures 5 B-C**), indicating specific recognition of antigens originating from the CHNB tumors (Passoni et al., 2002, Pistoia et al., 2012). These results suggest that the expression of tumor specific epitopes in NBs promotes immune recognition by the host T cells.

**Figure 5.**
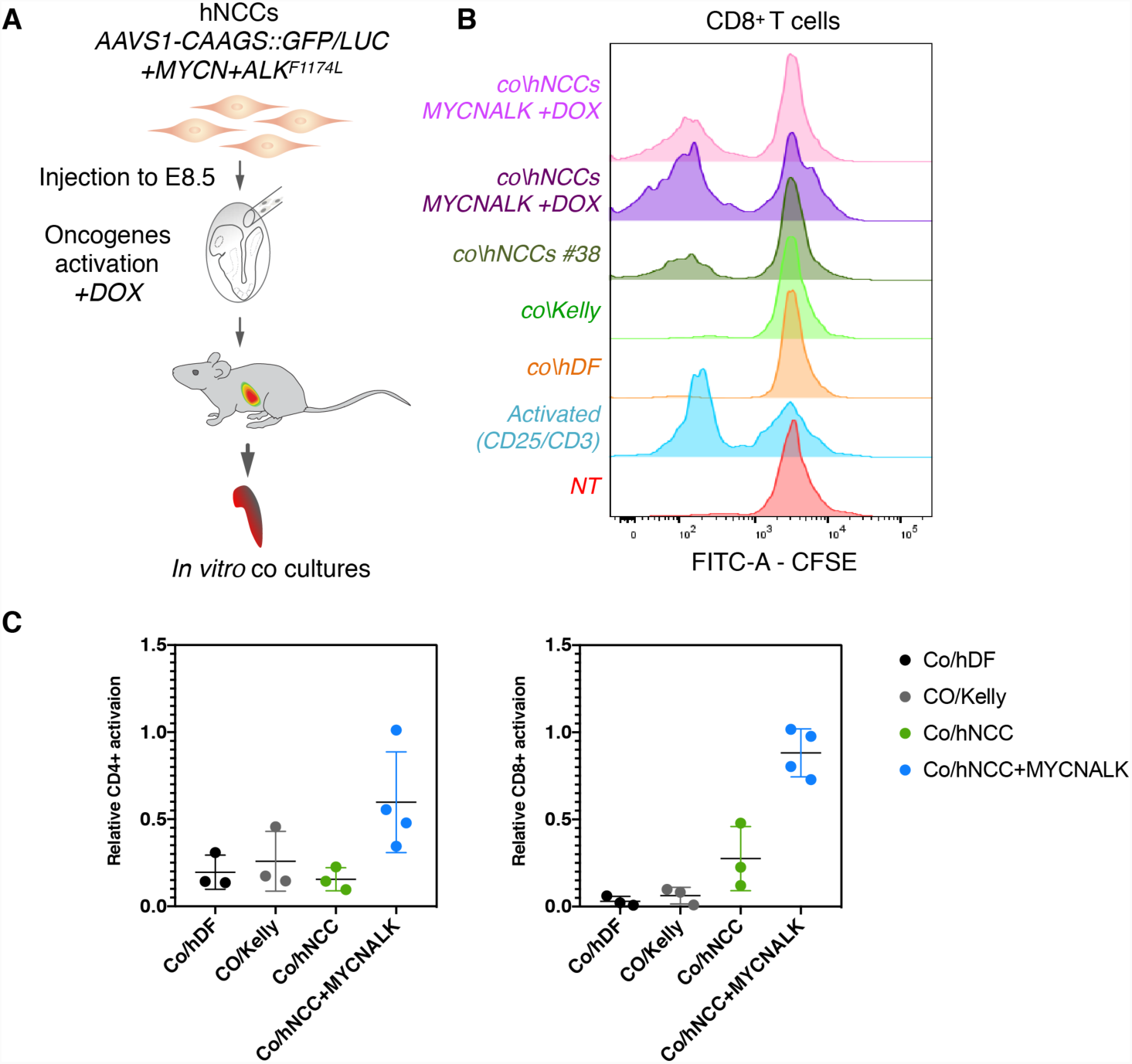
T Cells of NB Chimeric Mice Specifically Recognize Human Cells Originating in Human NB Tumors. (**A**) Schematic representation of the experiment. CFSE labeled splenocytes of chimeric mice were co-cultured *in vitro* in the presence of the hNCCs expressing or not expressing the ALK^F1174L^ and MYCN oncogenes, of human dermal fibroblasts (hDF; for HLA-I control) and of Kelly (NB cell line). T cell proliferation was monitored after 7 days. (**B**) CD8^+^ T cells of chimeric mice specifically recognized and proliferated when exposed to oncogene-activated hNCCs, which were used to generate the NB chimeras but much less when stimulated by unrelated human NB cell lines or hNCC not expressing the oncogenes. Anti CD3 and CD28 antibodies were used to stimulate T cell as a positive control (Activated), and non-treated splenocytes as negative controls (NT). (**C**) Summary of multiple experiments presenting the proliferation CD4^+^ (right) and CD8^+^ (left) T cells subpopulations after co-cultures in vitro at different conditions. Proliferation rates were normalized to controls. Data presented as means, error bars represent SD.

## Discussion

The results described in this paper show that the human-mouse chimeric platform described here allows to model human neuroblastoma in an animal system. Oncogene induced tumors were heterogenous consisting of multiple cell types similar to primary patient derived NBs. This is in contrast to tumors induced in xenotransplantation models, which are homogenous and are lacking the expression of NB markers typically seen in patients. The tumors developed in immune-competent mice despite a vigorous host anti-tumor response as indicated by massive tumor infiltration of the tumors by CD8^+^ cytotoxic T cells. The immune response was blunted by recruitment of Tregs and the activation of check point inhibitors such as mouse Tim-3 and human PD-L1 allowing progressive tumor growth.

A characteristic feature of NB is its clinical heterogeneity, ranging from spontaneous to highly metastatic and poor prognosis disease, characterized by unfavorable biological hallmarks. The mechanisms underlying such disparate behavior are unclear, although recent evidence suggests that host immunity plays an important role in the evolution of NBs during development (Brodeur and Bagatell, 2014). While anti-GD2 antibody therapy has provided some success against NB (Sait and Modak, 2017), this approach and other immune-based strategies remain suboptimal due to toxicity and, more importantly, the rapid emergence of relapse. Previous studies show that MYC plays a key role in suppressing immune surveillance in human leukemias and lymphomas (Casey et al., 2016). Similarly, it was suggested that MYCN is also associated with repressing cellular immunity in NB (Zhang et al., 2017, Wei et al., 2018), and such immune tolerance has emerged as a hallmark for NB prognosis. Current transgenic mouse models for NB offer limited insights into the tumor immune microenvironment. It is possible that the intermediate expression level of MYCN in the chimeric model captures the dynamics of cancer development along with the relevant immune action and the tumor evasion response. Hence, our chimeric model provides a unique system for the study of NB development in the context of the MYCN-controlled immune evasion mechanisms. This may help understanding the mechanisms adopted by NB for immune evasion and will be beneficial for the development of future immunotherapeutic strategies.

Interspecies chimeric mice may develop thymic-tolerance to human donor cells during embryonic development of the chimeric hosts when exposed to the human NC cells. Indeed, in a similarly chimeric system reported by others, induction of central immune tolerance to human donor cells was observed in immunocompetent mice following injection of human colorectal cancer cells into mouse blastocysts (Chen et al., 2015). Further investigation is needed to determine whether this thymic-tolerance is achieved by direct contribution of human cells to the mouse thymic-development or by cross-antigen presentation of human antigens by dendritic cells in the developing thymus of chimeras. While the thymus epithelium was suggested to originate from the NC, there is no clear evidence for the involvement of NC derived cells in the T cell education processes within the thymus (Foster et al., 2008). Hence, the nature of interspecies tolerance in post-implantation chimeric systems remains to be investigated.

Our results show a clear anti-tumor immune response of the host against human NB tumors. This normally involves infiltration of immune cells, activation of IFN-γ signaling and of genes associated with an effective inflammation and immune response (Spranger, 2016). CD8^+^ T lymphocytes play a key role in immunity to cancer, within their capacity to kill tumor cells upon recognition by T-cell receptor of specific antigenic peptides presented on the surface of target cells. Our data show that the host CD8^+^ T cells infiltrate the human NB tumors and specifically react against oncogene related antigens, when co-cultured with relevant cells. This may suggest that specific tumor-associated antigens produced by the human donor hNCCs elicit recognition by sub-clones of T cells. A typical productive cytotoxic T cell infiltrate is accompanied by the upregulation of PD-L1 on the cancer cells (Spranger et al., 2013), similarly to our observation in the chimeric model. The immune reaction against CHNBs resembles the characteristic and typical anti-tumor immunity seen in patients with a common tumor-microenvironment signature arguing against xeno-rejection mechanisms being responsible for the immune response.

In summary, the NB-bearing mouse-human chimera platform provides a highly defined model for studying the initiation and progression of NB, as well as its tumor immune-microenvironment and the immune–tumor interactions which govern immune evasion. Importantly, the ability to grow clinically derived human NB tumor cells in immune-competent hosts will facilitate the study of critical interactions between high-risk NB and the immune system. Ultimately, the chimeric model system may prove to be an optimal platform for studying NB pathogenesis and enabling the evaluation of different combinations of anti-oncogenic and immune-based therapies.

## Acknowledgments

We thank F. Soldner, Y. Stelzer, X.S. Liu for advice; W. Salmon (W. M. Keck Biological Imaging Facility, WI), S. Gupta (Genome core, WI), K. Cormier (Hope Babette Tang Histology Facility, KI, MIT), W. Huang (Animal Imaging & Preclinical Testing Facility, KI, MIT), R. Alagappan, C. Garrett-Engele, S. Markoulaki, R. Flannery and D. Fu for experimental assistance. We thank Robert Weinberg (Whitehead Institute), Jianzhu Chen and Tyler Jacks (The Koch Institute for Integrative Cancer Research at MIT) labs for advice and materials. This work was supported by grants from the Emerald Foundation (R.J., M.A.C.), the LEO Foundation (L18015; R.J., M.A.C.), the St. Baldrick’s Foundation (R.E.G., R.J., S.S., M.A.C.) and by the R37HD045022, 1R01-NS088538 and 5R01-MH104610 NIH grants (R.J.).

## Author Contributions

M.A.C. and R.J. conceived this study. R.J. performed the initial *in-utero* injections. M.A.C. designed, preformed and analyzed all experiments. S.Z. assisted in preforming experiments and analyzing data. Sa.S., H.M., B.H., R.E.G. and St.S, provided experimental help, materials and advice. G.B preformed bioinformatic analysis. M.A.C and R.J. wrote the manuscript, with input from Sa.S., R.E.G and St.S.

## Materials and Methods

### Mice

C57BL/6, NSG, and W^sh^/W^sh^ mice were obtained from the Jackson Laboratory. CD1-Elite mice were obtained from Charles River Laboratories. Mice were maintained in the Whitehead Institute for Biomedical Research animal facility. All experiments were approved by the Committee on Animal Care (CAC) and the Department of Comparative Medicine (DCM) at the Massachusetts Institute of Technology (MIT) and animal procedures were performed following the National Institute of Health (NIH) guidelines.

### hPSCs culture and differentiation to hNCCs

The hESC line WIBR#3 (X,X; Lengner et al., 2010) and hiPSC line AA#1 (X,X; Cohen at al., 2016) were cultured as previously described (Lengner et al., 2010). Differentiation to NCCs was performed as previously described (Cohen at al., 2016). Briefly, cells were cultured in KOSR medium (Thermo Fisher Scientific) supplemented with 10 μM SB431542 and 500 nM LDN193189 and 3 μM CHIR99021 (all from Stemgent) for the first 72 hour after plating hESCs/hiPSCs at a density of 2-3 x10^5^ cells per cm^2^. Subsequently, a gradual switch from KOSR to N2/B27 medium (Thermo Fisher Scientific), supplemented with 3 μM CHIR99021 was used until day 11. At day 12 cell cultures were enriched with hNCCs and cultured in N2B27 supplemented with bFGF and EGF (both 20ng/ml, Peprotech) and 3 μM CHIR99021.

### Embryonic hNCCs microinjection

Microinjections of hNCCs were performed as previously described (Cohen at al., 2016). Briefly, pregnant females at E8.5 were anesthetized via an intraperitoneal injection of avertin. Laparotomy was performed and the uterus was exposed and held with forceps during the injection of each embryo. NCCs were drawn into a glass micropipette, and the tip of the glass micropipette was inserted into the distal third of the decidual swelling. Approximately 2-5 x 10^3^ cells (suspended in 0.2-0.8μl of cell culture media) were injected per embryo.

### Neural crest cell contribution

Pregnant female mice were sacrificed between E14.5 to E15.5 of gestation following institutional guidelines. Embryos were harvested via dissection from the uterus and the placenta. NCC contribution to the embryos was determined by the presence of a fluorescent protein signal. Embryos containing a fluorescent signal were imaged using a stereomicroscope (Nikon SMZ18).

### Oncogene activation

*ALK*^*F1174L*^ and *MYCN* genes were cloned into a lentiviral backbone under the doxyclycline (Dox)-inducible Tet-ON (TetO) promoter. hESCs or hiPSCs carrying the tetracycline transactivator (M2rtTA) were infected with virus encoding *ALK*^*F1174L*^ and *MYCN* transgenes followed by Dox administration to activate the oncogenes (2 µg/ml in culture media; 2 mg/ml in mouse drinking water; MilliporeSigma).

### Xenograft assay

1.5×10^6^ hNCCs resuspended in 200 μl Matrigel/DMEM/F12 (50%, *v/v*) were injected into the flanks of NSG mice. To activate the oncogenes, Dox was administrated in drinking water (2 mg/ml). Only immunocompromised mice, which had been injected with hNCCs overexpressing *ALK*^*F1174L*^ and *MYCN* transgenes and treated with Dox, developed xenograft tumors. Tumors were collected when reached the size of 1 cm^2^ (Mean = 9.1 ± 2.2 weeks; n=18)

### Monitoring tumor growth by optical imaging

For luciferase imaging mice were anesthetized with Isofluorane (1.5-2.0 %) and injected intaperitoneally with D-luciferin (30 mg/ml, Perkin Elmer), 5 µl/gr of body weight, using a 26-gauge needle. Mice were imaged using Caliper IVIS Spectrum.

### H&E (Hematoxylin and Eosin) IF (immunofluorescence) and IHC (immunohistochemistry) staining

Cells, tumors or tissues were collected and fixed in 4% paraformaldehyde in PBS (MilliporeSigma). Patient derived NB tissue microarray were obtained from US Biomax. For IHC, samples were embedded in paraffin and sectioned into 4μm/slide. Following deparaffinizing in xylene and alcohol gradient, antigen retrieval was performed in sodium citrate buffer. For immunostainings, samples were blocked with 2% BSA and incubated with primary antibodies including Rabbit anti-ALK (1:200, Cell Signaling), Rabbit anti-MYCN (1:50, Cell Signaling), Rabbit anti-CD3 (1:300, Thermo Fisher Scientific), Rabbit anti-CD8a (1:300, Cell Signaling), Rabbit anti-FoxP3 (1:75, R&D systems), Mouse anti-Il2ra (CD25, 1:100, Novus), Rabbit anti-human-PD-L1 (1:300, Cell Signaling), Sheep anti-human-CD47 (1:100, R&D systems), Rat anti-mouse F4/80 (1:100, Thermo Fisher Scientific), Rabbit anti-γH2AX (1:300, Abcam), Rabbit anti-human-Ki67 (1:20, Thermo Fisher Scientific), Rabbit anti-Chromogranin A (1:500, Novus), Rabbit anti-Synaptophysin (1:200, Cell Signaling), Mouse anti-human-Nestin (1:300, Abcam), Rabbit anti-TH (1:500, PelFreez), Rabbit anti-Peripherin (1:500, Abcam), Rabbit anti-human neurofilament (1:100, 160kD, Abcam) and anti-eGFP (1:1000, Aves Labs) overnight at 4°C followed by appropriate secondary antibody incubation for 1-2h (Thermo Fisher Scientific). IF samples were imaged using Zeiss LSM 710 confocal microscope. For H&E staining, deparaffinized sections were stained in hematoxylin, incubated with 1% acid alcohol (1% HCl in 70% EtOH) and ammonia water, and stained with 1% Eosin.

### Splenocyte isolation and in vitro co-culture

Splenocytes were collected from the chimeric mice. At a density of 5×10^6^ cells /mL the splenocytes were labeled with CFSE (2μM, Invitrogen) and plated in 96-well U-bottom plates. To activate T cells in vitro, 96-well U-bottom plates were coated with CD3 (1:100, BioLegend) and CD28 (1:100, BioLegend), and incubated at 37°C for 2h. For co-culture, mitotically inactivated (mitomycin-C, 10μg/ml for 2 hours, MilliporeSigma) human dermal fibroblasts (hDF), Kelly cells or hNCCs were plated with the splenocytes in a 1:10 ration. Cultures were maintained at 37°C, 5% CO_2_ in complete RPMI, supplemented with β-ME (50μM; MilliporeSigma) and HEPES (10mM; Thermo Fisher Scientific). After 7 days of incubation, cells were collected and analyzed by flow cytometer.

### Flow cytometry

Single-cell suspensions were prepared and splenocytes were incubated with APC-conjugated anti-CD4 (0.25μg/10^6^ cells in 100μl volume, BioLegend) and Perp-Cy5.5-conjugated anti-CD8 (1μg/10^6^ cells in 100μl volume, BioLegend) for 30 min on ice. For HLA class I analysis, total cell number was counted, and 2×10^6^ cells were incubated with mouse anti-HLA class I antibody (1:200, BioLegend) for 30min followed by secondary antibody incubation for 30min. Cells were analyzed by flow cytometry on LSRII or LSRFortessa device with BDFacs DIVA software. The data collected was analyzed by Flow Jo software (TreeStar).

### RNA-seq

Cells and tumor samples were homogenized (QIAshredder), and RNA was extracted (RNeasy, Qiagene). Poly(A) mRNA capture and construction of stranded mRNA-Seq libraries were made using KAPA mRNA HyperPrep Kit according to instructions (Roche Sequencing Solutions). NGS was performed at the Whitehead Genome Technology Core (HiSeq 2500, Illumina). To distinguish reads derived from human and mouse cells, we created a metagenome of human (GRCh38) and mouse (GRCm38) canonical chromosomes and a matching metatranscriptome annotation (GTF) file, containing Ensembl Release 91 gene models. This metagenome was indexed for STAR, using the gtf file with “--sjdbOverhang 39”, and reads were mapped with STAR using default parameters. Gene counts were obtained using featureCounts (with “-s 2”) and subsequently normalized with DESeq2. For comparison with public RNAseq datasets (with gene counts expressed as FPKM), gene counts were obtained with cufflinks, using NCBI RefSeq gene models (downloaded from UCSC Bioinformatics on 4 Oct 2017).

### Quantitative Image Analysis

IHC images were processed into separate channels representing nuclei staining (haematoxylin) and IHC staining (DAB) using Fiji. Images were then analyzed using the open-source CellProfiler software (www.cellprofiler.org; Kamentsky et al., 2011) for cell segmentation and staining characterization. Briefly, cell nuclei and IHC staining were segmented based on Gaussian blurred images in both haematoxylin and DAB channels respectively using size and intensity thresholds. Cells were then filtered based on the channels overlapping rate, and those which exhibit over 60% overlapping rates, representing cells expressing the tested markers, were scored and quantified. The software pipeline detection was tested to be highly correlated with visual detection.

### LacZ staining

Cell were fixed shortly with 2% formaldehyde and 0.5% glutaraldehyde in PBS, and incubated in staining-solution containing 1 mg/ml X-Gal, 5 mM potassium ferricyanide, 5 mM potassium ferrocyanide and 2mM MgCl2, in PBS (All from MilliporeSigma) at 37^0^C, over-night.

### Southern Blot

10µg of DNA isolated from hPSCs was digested overnight with the appropriate enzymes and run on a 0.8% agarose gel. DNA was transferred to a nylon membrane (Amersham) and hybridized with 32P-labeled random primer (Stratagene) probes.

### Quantification and Statistical Analysis

Statistical parameters including the exact value of n and measures (mean ± SD) and statistical significance are reported in the Figures and/or the Figure Legends. Data are judged to be statistically significant when P < 0.05 by two-tailed Student’s T-Test or by 2-way ANOVA, where appropriate.

### Western Blot

Cells were lysed by RIPA buffer with proteinase inhibitor (Invitrogen), and subject to standard immunoblotting analysis. Rabbit anti-ALK (D5F3 and C26G7), anti-p-ALK (Y1604), anti-MYCN and anti-p44/42 MAPK (Erk1/2) antibodies were used (Cell Signaling).

**Figure S1.**
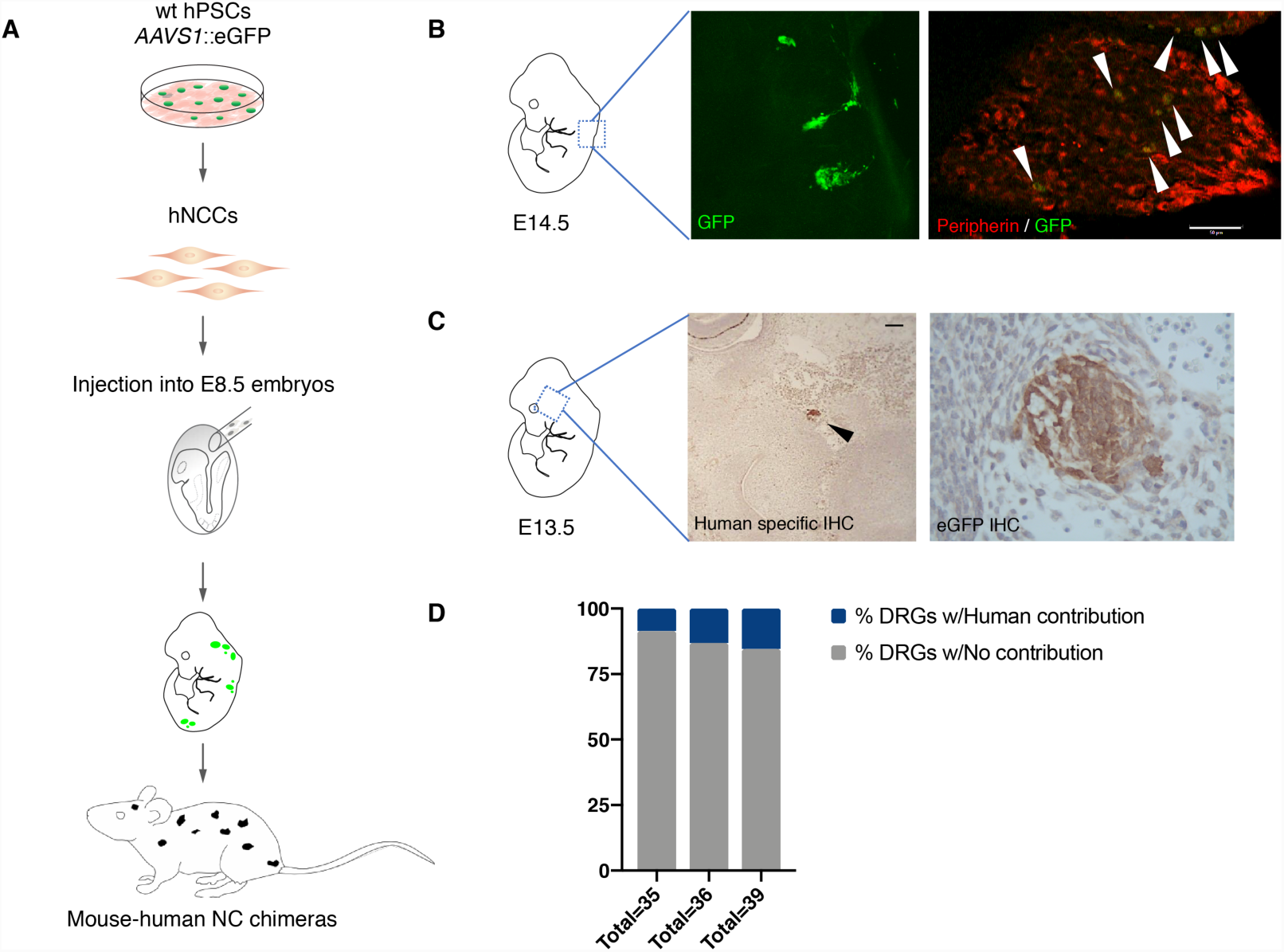
Human NCCs Contribute to PNS Subsets in Interspecies Chimeras After in Utero Injection Into Mouse Embryos. (**A**) *AAVS1-CAAGS::eGFP* hPSC-derived hNCCs contribute to PNS embryonic development as indicated by the detection of eGFP in the embryos. (**B**) eGFP positive human cells were found in structures resembling dorsal root ganglia (DRG) in an E14.5 mouse embryo (**B**, Left). Contribution of human cells to DRGs was confirmed by immunofluorescence staining for eGFP (human donor cells) and for Peripherin, a PNS marker (**B**, right; double positive cells are indicated by arrowheads; scale bar=50µm). (**C**) Human cells were found to contribute to the trigeminal ganglion of an E13.5 mouse embryo. Contribution of human cells was confirmed by immunohistochemistry (IHC) using anti-human specific antibody (**C**, left; black arrowhead; scale bar=100µm), and anti eGFP antibody (**C**, right). (**D**) hNCCs were found to contribute to DRGs of adult chimeric mice. Single DRGs of chimeric mice were collected and analyzed for human contribution by a qPCR assay (Cohen at al., 2016). Three representative chimeric mice present chimeric contribution in 8-15% of tested DRGs. The total number of tested DRGs per mouse is indicated.

**Figure S2.**
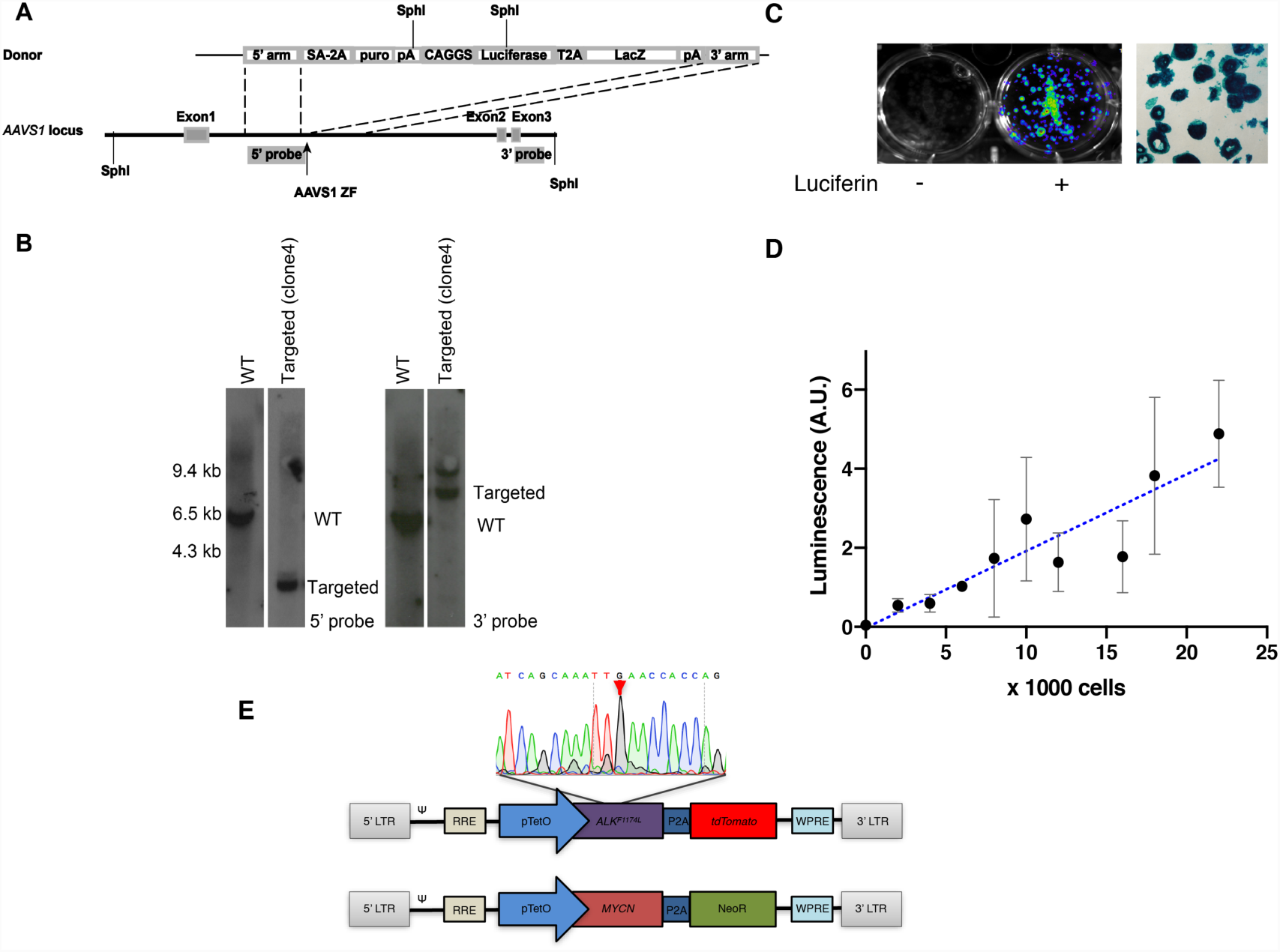
Generating Luciferase Expressing Donor hESCs for Non-Invasive *in Vivo* Imaging. (**A**) Targeting strategy to generate donor human cells with the *CAGGS-Luciferase-t2A-LacZ* cassette integrated to the *AAVS1* locus. (**B**) Southern blots of parental WIBR#3 hESCs (WT) and homozygote-targeted cells (clone #4) with the 5’ internal probe (left panel) and 3’ external probe (right panel). (**C**) Targeted WIBR#3 clone #4 hESC cultures show Luciferase expression upon Luciferin treatment (left panel), and targeted hESC colonies show LacZ expression, demonstrated by X-gal staining (right panel). (**D**) Quantification of luminescence activities of targeted cells show a linear correlation of the luminescence levels within a range of cell numbers of hESCs cultures (0-22K cells), demonstrating a highly sensitive assay for detection of human cells (Linear regression *p*-value <0.0001; Data presented as means of triplicate reads, error bars represent SD). (**E**) A schematic representation of the *FUW-TetO-ALK*^*F1174L*^*-p2A-tdTomato (*top) and *FUW-TetO-MYCN-p2A-NeoR (*bottom) lentiviral backbones made for *ALK*^*F1174L*^ and MYCN overexpression. Sanger sequencing confirmed the C to G transition leading to the *ALK*^*F1174L*^ specific mutation (arrowhead).

**Figure S3.**
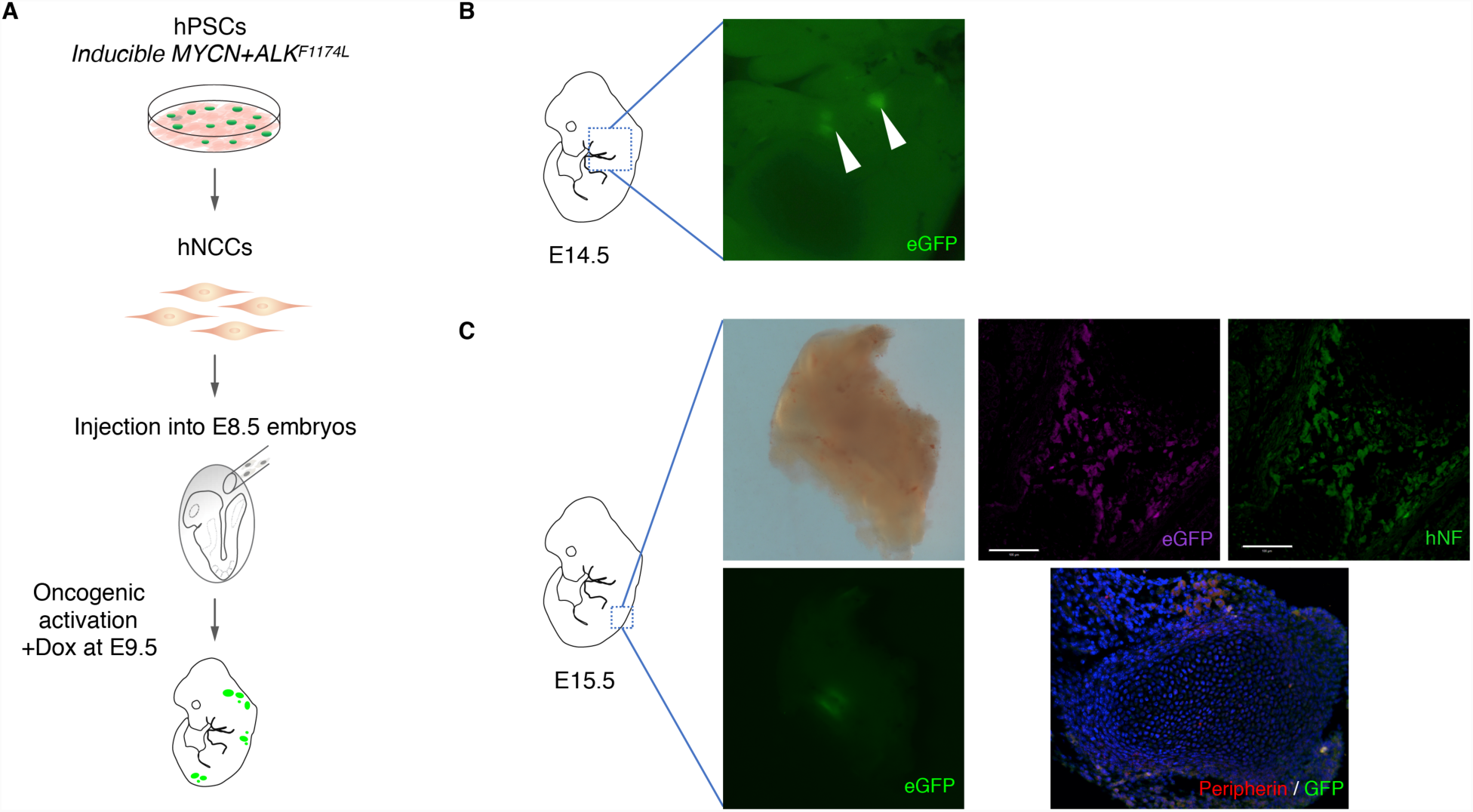
hNCCs Contribute and Proliferate in Internal Structures of Host Embryos. (**A**) A schematic representation of the experiment. hPSCs were differentiated into NCCs and were microinjected into E8.5 mouse embryos to generate mouse-human NC chimeric embryos. Oncogene activation was started at E9.5 using Dox. (**B**-**C**) Representative embryos, injected with hNCCs and evaluated for human cells at E14.5-15.5. hNCCs contribute to embryonic development and form aggregates, visible by eGFP in the trunk (**B**) or abdomen (**C**) area, as an indicator for donor cell proliferations *in vivo*. (**C**) Cell aggregates of donor cells were immunostained for human specific markers (upper panel; anti-eGFP in magenta and anti-human neurofilament, hNF, in green) and differentiation marker (lower panel; anti-Peripherin in green and anti-eGFP in red) to confirm their human origin and their differentiation capacity *in vivo*.

**Figure S4.**
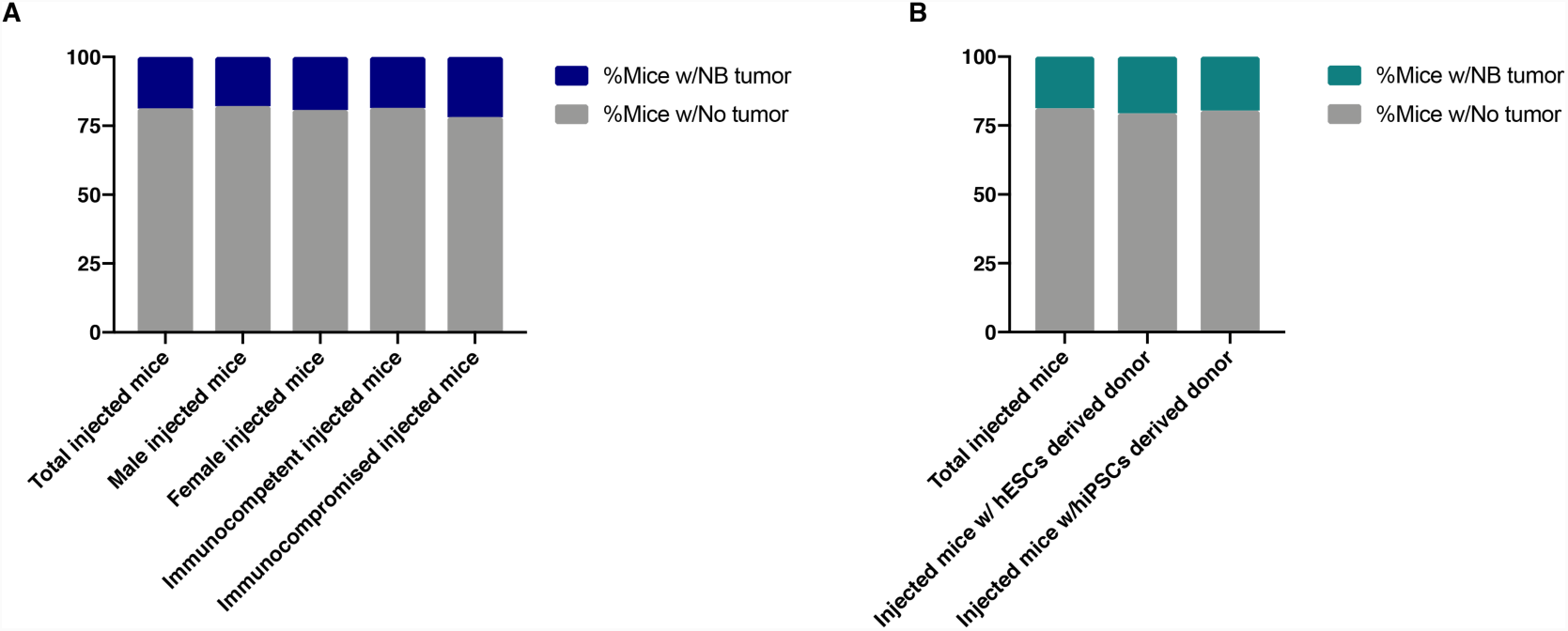
Summary of Human NB Development in Chimeric Mice Injected *in Utero* With Oncogene Expressing hNCCs. Out of a total of 144 adult mice which were injected at E8.5, 27 developed human NB tumors. (**A**) The distribution of chimeric mice which developed tumors was similar when human cells were injected into male (n=56) and female (n=88) hosts, and when injected into immunocompetent (n=108) or immunocompromised (n=36) hosts. (**B**) hNCCs donor cells derived from two different hPSC lines (hESCs: n=41; hiPSCs: n=103) with different genetic background produced similar rates of NB tumor development.

**Figure S5.**
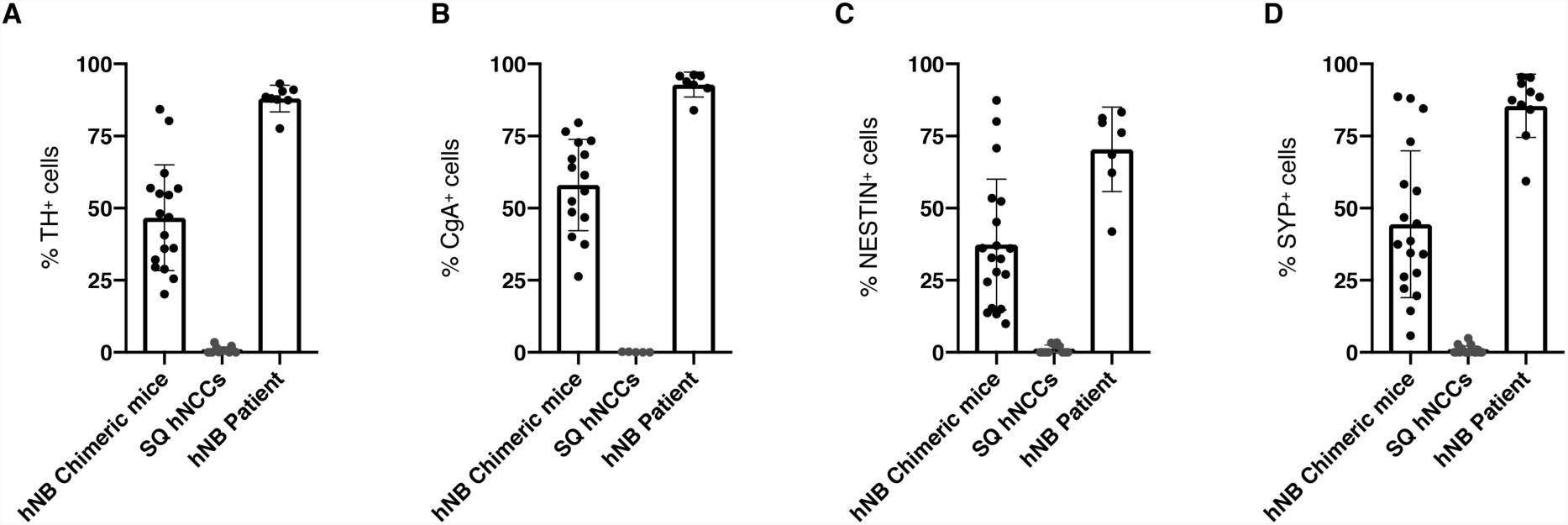
Tumor of Mouse-Human NC Chimeras Express Markers of NB. Quantifications of IHC Experiments for the Percentage of Positive Cells Expressing the Typical NB Markers. (**A**) Tyrosine hydroxylase (TH), (**B**) Chromogranin A (CgA), (**C**) Nestin (NES), and (**D**) Synaptophysin (SYP), in samples of human NBs from chimeric mice (n=5), subcutaneous xenograft outgrowth of hNCCs (n=4), and NB samples of patients (Data presented as means, error bars represent SD, dots represent fields of view).

**Figure S6.**
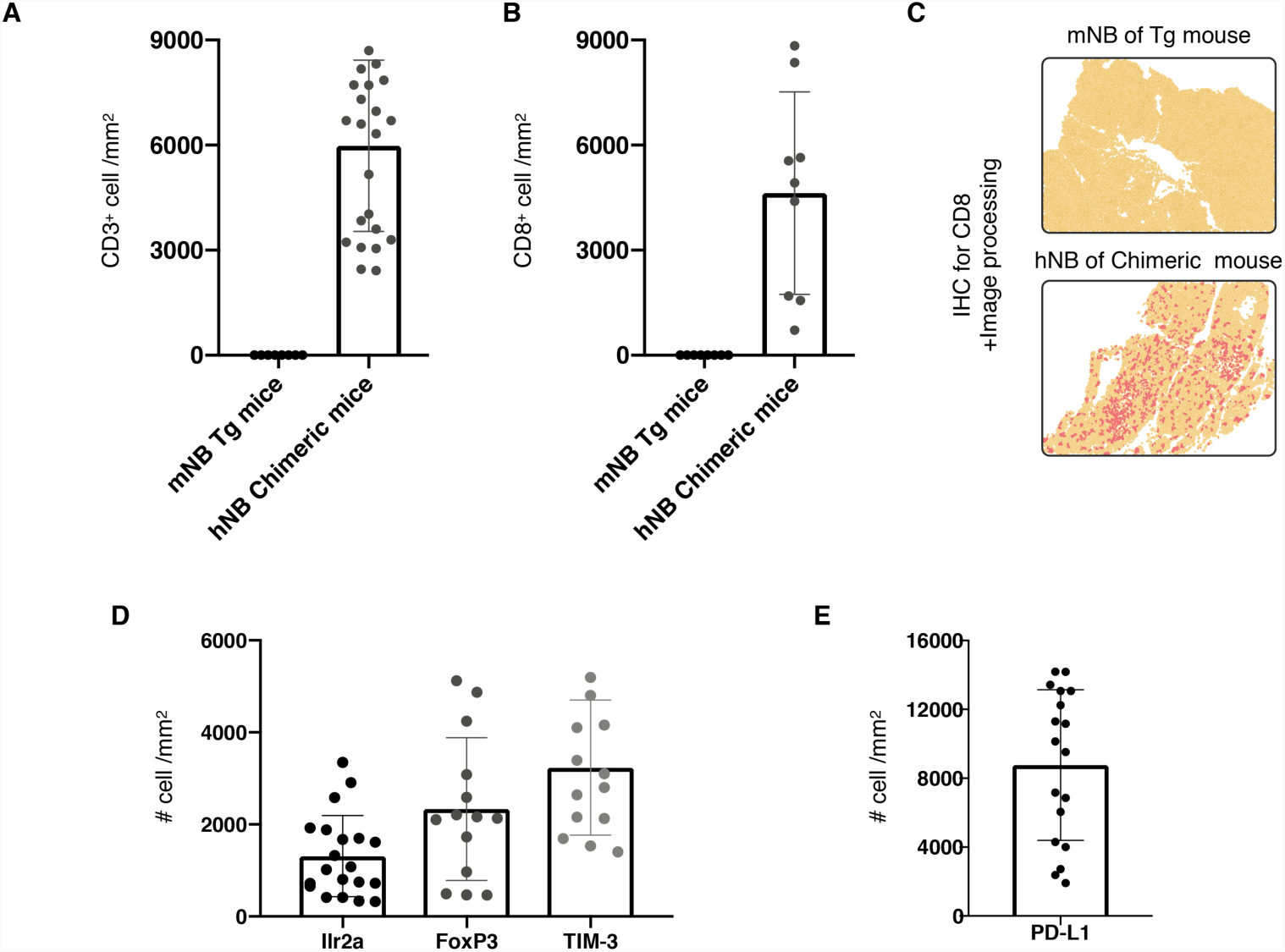
NB Tumor of Mouse-Human Neural Crest Chimeras Are Highly Inflamed and Infiltrated by Host T Cells, Tregs and Activate Checkpoint Markers. Quantifications of IHC experiments for mouse specific CD3 (**A**) and CD8a (**B**) by comparing tumors of mouse transgenic models of NB (Berry et al., 2012; n=2) and human NB in chimeric mice (n=6; Tg: transgenic). The rates of infiltrating cells were measured and quantified by image analysis for immune score (Galon et al., 2006; See material and methods). (**C**) Representative fields of IHC for CD8a following software analysis (Yellow=tissue area, Dark yellow=nuclei, Red=DAB staining of CD8a IHC). (**D**) Quantifications of IHC experiments for mouse specific Ilr2a, FoxP3 and Tim-3 and (**E**) human specific PD-L1 (n=5; Data presented as means, error bars represent SD, dots represent fields of view).

**Figure S7.**
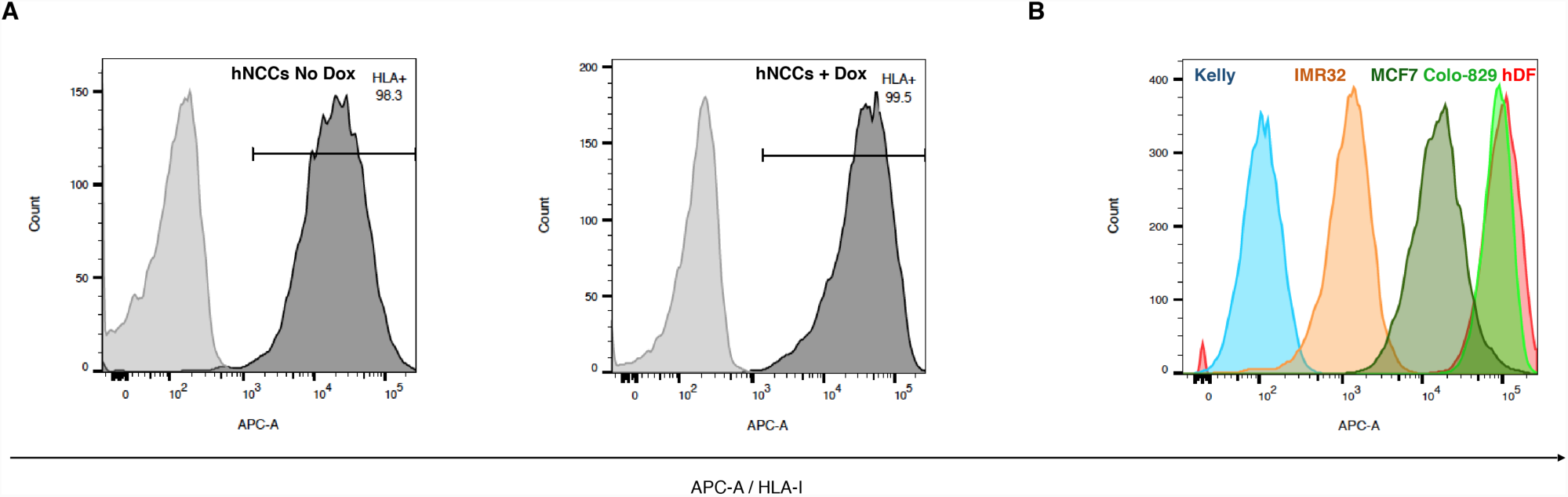
The Expression Level of HLA-I in Human Cells. To Measure the Expression Levels of HLA-I, FACS Analysis Was Performed. (**A**) A representative FACS assay for hPSC-derived hNCCs using HLA-I antibody (Dark grey) and control antibody (Light grey) show that over 98% of hNCCs express HLA-I, both, with (left panel; +Dox) or without (right panel; No Dox) the over-expression of MYCN and ALK^F1174^. (**B**) A representative FACS assay for HLA-I for a panel of human cells demonstrate a graduate expression levels of HLA-I: While Kelly (Blue) and IMR-32 (Brown), both NB cell lines, minimally express HLA-I, the MCF7 (Dark Green; Brest cancer), Colo-829 (Light green; Melanoma), and primary human dermal fibroblasts (hDF, in red) express higher level of HLA-I.

